# Distinct subcortical pathways mediate collision detection with and without attention and consciousness

**DOI:** 10.64898/2026.01.16.700036

**Authors:** Jinyou Zou, Ye Wang, Peng Zhang

## Abstract

Accurate detection of impending collisions is essential for survival. While multiple subcortical pathways have been found in the detection of looming stimuli, the roles of top-down attention and consciousness in this process remains unclear. Using high-resolution fMRI, we investigated how top-down attention and conscious awareness influence the subcortical processing of collision trajectories in healthy participants and hemianopic patients with cortical blindness. In healthy participants, the superior colliculus (SC)-parabigeminal nucleus (PBGN)-amygdala pathway integrated binocular information for accurate detection of collision trajectories in the upper visual field, but only when participants were paying attention to the looming stimuli. On the other hand, for stimuli presented in the blind visual field of hemianopic patients, effective connectivity analyses revealed collision sensitivity in the SC-pulvinar and SC- ventral tegmental area (VTA) pathways, supporting automatic collision detection even in the absence of visual awareness. Together, these findings suggest that distinct subcortical pathways mediate collision detection in the human brain depending on the presence and absence of attention and conscious awareness.

**Highlights:**

- Attention selectively enhanced collision sensitivity in the SCs-PBGN-CMA pathway.
- Binocular information contributed to accurate collision detection in the upper visual field.
- SC-Pulvinar and SC-VTA pathways detect collision trajectory without visual awareness.

**Significance statement:** Accurate detection of collision trajectory is critical for survival. Using high-resolution fMRI in healthy participants and hemianopic patients, we show that collision detection is mediated by distinct subcortical circuits depending on the attentional state and conscious awareness. The SC–PBGN–amygdala pathway can precisely detect collision trajectory in upper visual field with top-down attention, whereas the SC–pulvinar and SC–VTA pathways enable automatic collision detection without awareness. These findings reveal distinct functional roles of the subcortical pathways in collision detection.

## Introduction

The ability to detect impending collisions is vital for our survival. Subcortical mechanisms for processing looming information have been found in both animals (Wang and Frost, 1992; Sun and Frost, 1998; Shang et al., 2015; Salay et al., 2018) and humans (Billington et al., 2011; Guo et al., 2024; Thieu et al., 2024). In rodents, recent studies have identified multiple subcortical circuits from the superior colliculus (SC) that are involved in innate defensive responses to looming stimuli, including SC-PBGN (parabigeminal nucleus)-Amygdala (Shang et al., 2015), SC-LP (lateral posterior nucleus of the thalamus, a pulvinar-like structure in rodents)-Amygdala (Wei et al., 2015), and SC-VTA (ventral tegmental areas)-Amygdala (Zhou et al., 2019). In humans, the SC-VTA and SC-pulvinar pathways also exhibited collision sensitivity even without attention and awareness to the looming stimuli (Guo et al., 2024). However, it remains unclear whether and how top-down cognitive influences modulate collision detection mechanisms in these subcortical pathways (Tseng et al., 2023).

SC is a key structure in the mammalian brainstem involved in the control of attention and eye movement, integrating bottom-up sensory input and top-down signals from frontoparietal attention networks (May, 2006; Zhaoping, 2016). Optogenetic stimulation of the tectofugal pathways has been shown to trigger different types of defensive behaviors (Shang et al., 2018), depending on the internal and environmental context (Isa et al., 2020; Tseng et al., 2023), which may suggest that the SC and its downstream targets can be modulated by top-down influences. In humans, the SC and pulvinar exhibited stronger responses to looming compared to receding stimuli when performing a time-to-collision (TTC) judgement task (Billington et al., 2011). However, while human observers can precisely discriminate direct-hit versus near-miss trajectories (Lin et al., 2009; Duke and Rushton, 2012; Chen et al., 2016), SC activity showed no difference between these conditions during the trajectory discrimination task (Fig. 2c/2e in Guo et al. (2024)). Given the short TTC of looming stimuli used in this previous study, and the high sensitivity of SC activity to TTC, the absence of collision-sensitive activation may have resulted from response saturation in the SC. Response saturation may also explain the lack of collision sensitivity observed in the upper visual field, despite behavioral evidence showing an upper visual field advantage in collision detection performance (Guo et al., 2024). Thus, at longer TTCs, it is possible that top-down attention can enhance collision sensitivity in the SC to support accurate detection of collision trajectory, especially those from the upper visual field.

While retinal inputs from the two eyes are anatomically segregated in different layers and columns in the SC (Hubel et al., 1975; Dilbeck et al., 2022), SC neurons respond robustly to visual stimuli from both eyes (Cynader and Berman, 1972; Schiller et al., 1974). These findings suggest that the dendrites of SC neurons can integrate binocular information for visual processing, possibly including precise estimation of looming trajectories. In support of this, some SC neurons have preferred motion directions that differ by approximately 180 degrees between the two eyes (Tailby et al., 2012), consistent with a potential role in computing motion-in-depth. Moreover, behavioral evidence has shown that human observers can utilize binocular information for accurate detection of collision trajectories, exhibiting higher collision sensitivity to hit-nasion than hit-eye trajectories (Guo et al., 2024). These findings suggest that top-down modulation may enhance collision sensitivity in the SC via binocular integration of motion-in-depth information.

In this study, we aim to investigate how top-down attention and conscious awareness modulates collision-sensitive activations in human subcortical pathways originating from the SC, and whether subcortical activity reflects the upper visual field advantage and the use of binocular information in collision detection. To avoid response saturation observed in the previous study, we employed looming stimuli with a longer TTC in the fMRI experiment (138 ms vs. 33 ms in Guo et al. (2024)). In separate sessions, participants either attended to and judged the trajectories of the approaching objects (attended condition) or performed a demanding central fixation task (unattended condition). The results showed collision-sensitive responses to hit-nasion trajectories from the upper visual field in the superficial SC, PBGN, and centromedial amygdala in the attended condition. Path analysis of collision sensitivity further revealed significant SC–PBGN connectivity, specifically for stimuli in the upper visual field. In contrast, the SC-pulvinar and SC-VTA pathways support automatic collision detection without attention and awareness in the blind visual field of hemianopic patients.

## Results

The approaching object was a 3-D sphere presented on a 3-D screen, flying through a virtual tunnel towards the observer at a speed of 24 m/s. The sphere traveled from 11.3 m to 3.3 m in front of the observer (Fig. S1). The TTC of the last frame was 138 ms, much shorter compared to the previous study (33 ms), in order to avoid response saturation in subcortical regions. The trajectory would either hit the nasion (hit nasion), hit the ipsilateral eye (hit eye), or just missed the head (near miss) of participants (Fig. 1). In the attended condition, participants responded whether the approaching object would hit or miss their heads. In the unattended condition, they performed a demanding rapid serial visual presentation (RSVP) task at fixation and ignored the looming stimuli. To control for the effects of luminance-induced pupil dilation on subcortical responses, bright stimuli were presented on a black background in 11 participants, while dark stimuli were presented on a light gray background in another 12 participants. Checkerboard flickers in the same size and location as the looming stimuli were employed to localize retinotopic regions of interest (ROIs).

**Fig. 1.**
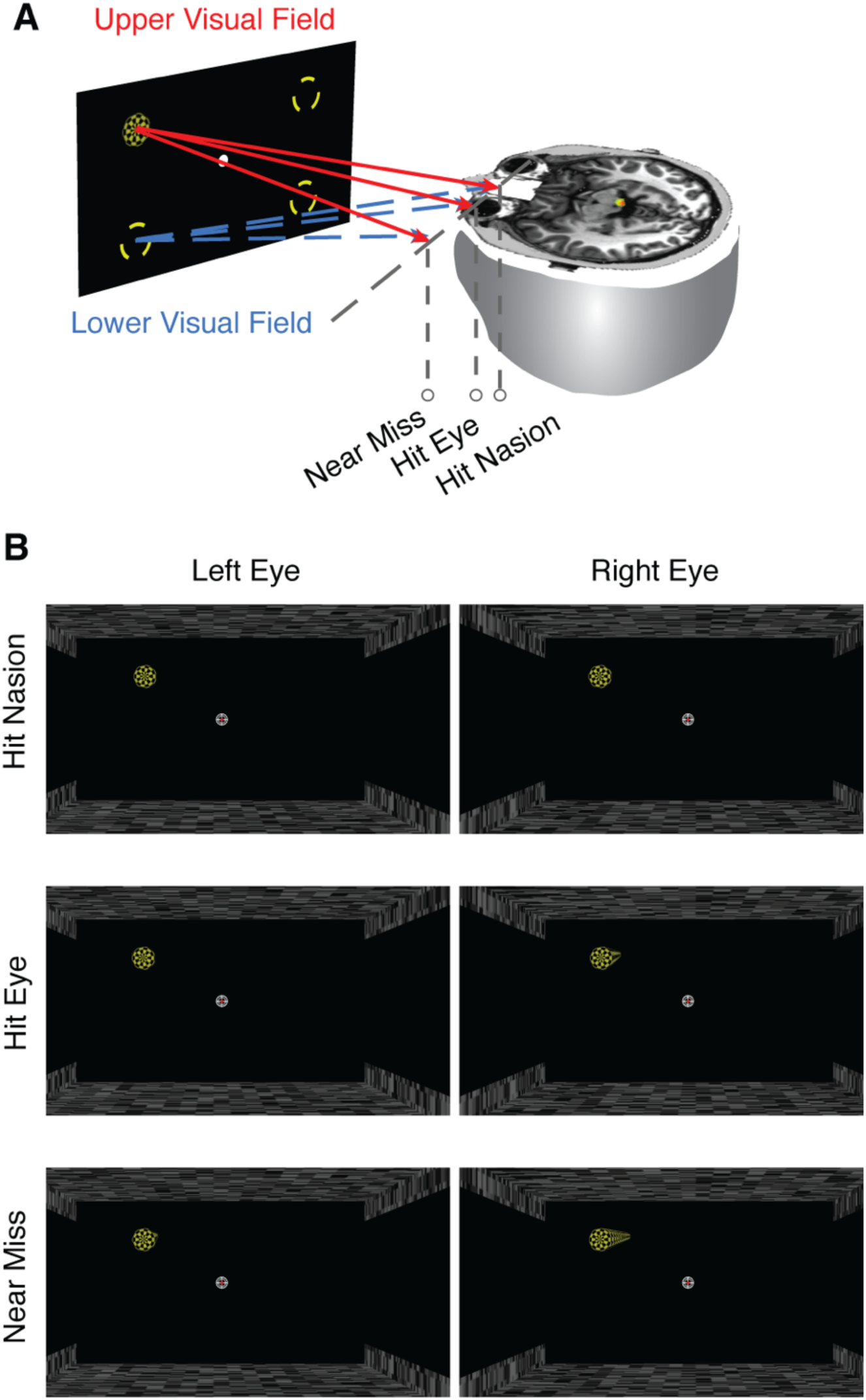
Schematic diagrams of looming stimuli. (A) Visual stimuli depicting an approaching object were presented using a MRI-compatible 3D monitor and polarized glasses. The trajectory would either be a direct hit (hit nasion, hit eye) or a near miss of the observer’s head. (B) The images presented to the left and right eyes were shown for three trajectory conditions. All video frames were superimposed to illustrate the motion trajectory.

## 1. Top-down attention enhanced UVF collision sensitivity in the superficial SC, PBGN and CMA

We first examined fMRI responses in the superior colliculus (SC). Given that neurons in the superficial and intermediate layers may be differently involved in processing looming stimuli (Shang et al., 2015; Wei et al., 2015), we analyzed layer-dependent responses in the SC. Based on the laminar organization of the human SC (Tardif and Clarke, 2002), voxels within 1.5 mm below the SC surface were defined as the superficial layers, while those between 1.5 mm and 3 mm in depth were classified as the intermediate layers. The retinotopic localizer predominantly activated voxels in the superficial layers, consistent with their role in processing visuosensory information (May, 2006; Zhang et al., 2015; Zhang et al., 2016). Thus, for the superficial SC, we selected voxels within the superficial layers that were significantly activated by the retinotopic localizer as the ROI. Separate ROIs were defined for the upper (UVF) and lower visual fields (LVF) (Fig. 2A). All voxels within the intermediate layers were included as the ROI for the intermediate SC.

**Fig. 2.**
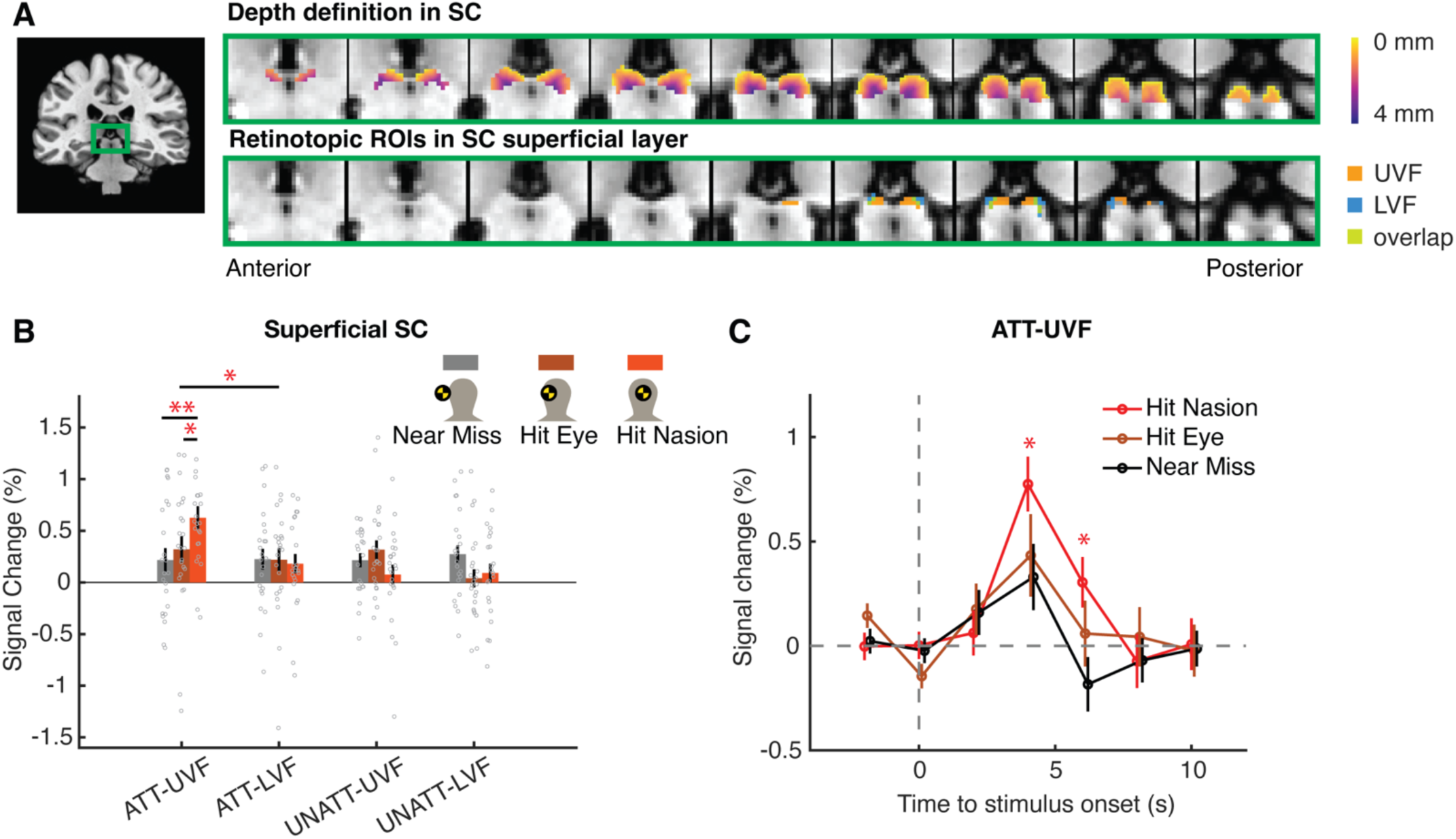
ROIs and fMRI responses of the superior colliculus (SC). (A) The upper and lower rows illustrate the depth map of SC voxels and the retinotopic ROIs in the superficial layers, respectively. Coronal slices were presented in a posterior-to-anterior sequence with a 1-mm step. (B) BOLD responses of the superficial SC to looming stimuli in different conditions. Dots denote the data of individual subjects. (C) Time courses of BOLD signals evoked by stimuli in the upper visual field under the attended condition. * p<0.05 (for hit-nasion vs. near-miss conditions in (C)), ** p<0.01. Error bars indicate S.E.M. ATT, attended; UNATT, unattended; UVF, upper visual filed; LVF, lower visual field.

A linear mixed effects (LME) analysis was conducted on ROI-averaged responses (Fig. 2B), separately for the attended (ATT) and unattended (UNATT) conditions. Trajectory (hit nasion / hit eye / near miss) and visual field (UVF / LVF) were included as fixed effects, with participant as a random grouping factor. In the attended condition, a significant interaction was observed between trajectory and visual field (F(2, 84) = 3.54, p = 0.034), along with a significant effect of trajectory in the UVF (F(2, 42) = 5.43, p = 0.008; hit nasion vs. near miss: t(22) = 3.18, p = 0.008, Cohen’s d = 0.756; hit nasion vs. hit eye: t(22) = 2.33, p = 0.049, d = 0.554), but not in the LVF (p = 0.932). In addition, stimulus luminance was included as a between-subject factor in the LME models and did not yield any significant effect (all p > 0.1), indicating that the observed collision sensitivity in the superficial SC was not attributable to changes in stimulus luminance. Notably, collision sensitivity (hit nasion vs. near miss) was robust across different voxel inclusion thresholds for the retinotopic ROIs (Fig. S2). Event-related average of time series (Fig. 2C) further revealed significantly greater responses to hit-nasion compared to near-miss trajectories in the ATT-UVF condition (peak response at 4s from stimulus onset: t(22) = 2.25, p = 0.035, d = 0.469; see Fig. S3 for other conditions). In the unattended condition, however, there were no significant main effects of trajectory (p = 0.158), visual field (p = 0.356), or their interaction (p = 0.11). In the intermediate SC, no significant effects of trajectory or interaction were found (all p > 0.1; Fig. S4).

The downstream targets of the SC include the PBGN (Shang et al., 2015), pulvinar (Wei et al., 2015), and VTA (Zhou et al., 2019). ROIs for the PBGN and VTA were defined using anatomical landmarks on the T1-weighted structural images (Fig. 3A) (Ding et al., 2016; Mai, 2016), whereas pulvinar ROIs were delineated based on retinotopic activations and a retinotopic atlas (Arcaro et al., 2015). In the attended condition, LME analysis of ROI-averaged responses (Fig. 3B) revealed a significant trajectory × visual field interaction in the PBGN (F(2, 84) = 3.91, p = 0.024), along with a significant effect of trajectory in the UVF (F(2, 42) = 5.92, p = 0.005; hit nasion vs. near miss: t(22) = 3.12, p = 0.01, d = 0.685; hit nasion vs. hit eye: t(22) = 2.81, p = 0.015, d = 0.618), but not in the LVF (p=0.761). No significant trajectory effects or interactions were observed in the pulvinar or VTA (all p > 0.38). In the unattended condition, no significant main effects of trajectory or trajectory × visual field interactions were found in any of the ROIs (all p > 0.13; Fig. 3C).

**Fig. 3.**
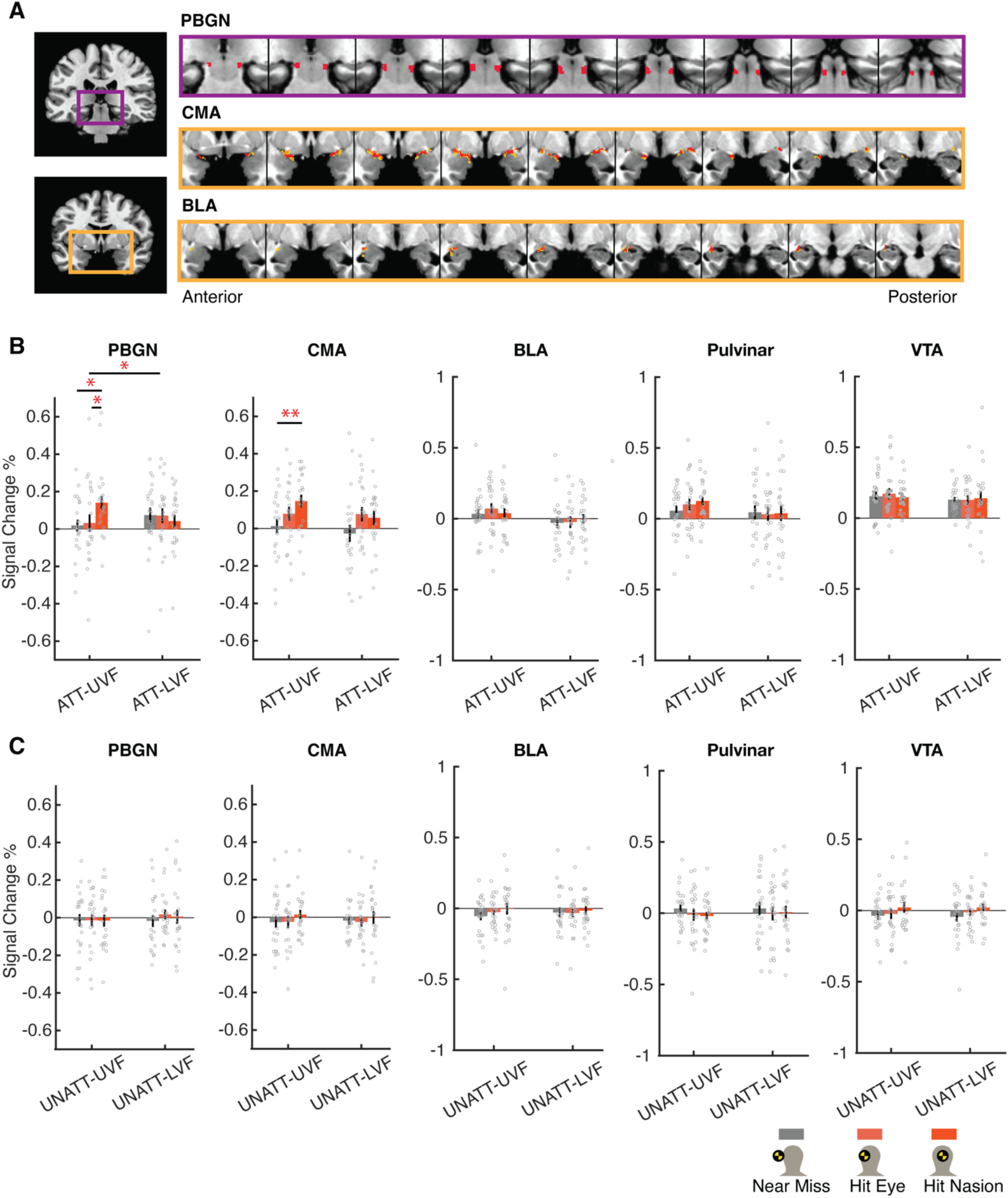
ROIs and fMRI responses of other looming-sensitive nuclei. (A) ROIs for PBGN and amygdala are illustrated with purple and orange frames, respectively. Coronal slices were presented in a posterior-to-anterior sequence with a 1-mm step. (B, C) BOLD responses of the downstream nuclei of SC to looming stimuli in the attended (B) and unattended (C) conditions. PBGN, parabigeminal nucleus; CMA, centromedial complex of amygdala; BLA, basolateral complex of amygdala; VTA, ventral tegmental areas. Other conventions as in Fig. 2.

Finally, we examined the amygdala, a key terminal of the tectofugal pathways and a common downstream target of the PBGN, pulvinar, and VTA (Shang et al., 2015; Wei et al., 2015; Zhou et al., 2019). The amygdala was divided into the centromedial (CMA) and basolateral (BLA) nuclei based on an anatomical atlas (Amunts et al., 2005; Eickhoff et al., 2005), and ROIs within each nucleus were further refined using functional connectivity with the superficial SC (Fig. 3A; see Methods for details). In the attended condition only (Fig. 3B), the CMA showed a significant main effect of trajectory (F(2,84) = 6.17, p = 0.003), which was driven by a trajectory effect in the UVF (F(2,42) = 5.12, p = 0.01; hit nasion vs. near miss: t(22) = 3.2, p = 0.008, d = 0.799), but not in the LVF (p = 0.106). No significant trajectory effects were observed in the BLA (all p > 0.496).

These findings showed that top-down attention selectively enhanced collision sensitivity (hit nasion vs. near miss) in the superficial SC, PBGN, and CMA, particularly for looming stimuli in the upper visual field.

## 2. Minimal Collision Sensitivity in the Geniculostriate Pathway and Higher Cortex

To investigate whether the collision sensitivity observed in the SC, PBGN, and CMA originated from the geniculostriate pathway and higher cortical regions, we examined fMRI responses to looming stimuli in the lateral geniculate nucleus (LGN) of the thalamus, early visual cortices, and frontoparietal regions. No significant effects of trajectory (all p > 0.1) were found in the LGN (Fig. S4), early visual cortices (Fig. S5), or frontoparietal regions (defined by the HCP-MMP1 atlas) (Fig. S5) (Glasser et al., 2016). To further examine collision sensitivity, we calculated a collision-sensitive response (CSR), defined as

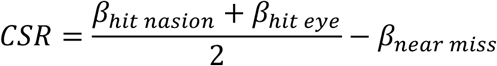

in which *β* indicated the fMRI responses in a certain condition. No significant CSR was observed in the ROIs of the early visual areas or other cortical areas in a whole-brain analysis (Fig. 4). Using hit nasion vs. near miss as the collision-sensitive response yielded similar results. These findings suggest that the collision sensitivity observed in the subcortical areas was unlikely originated from the geniculate-striate pathway or higher cortex.

**Fig. 4.**
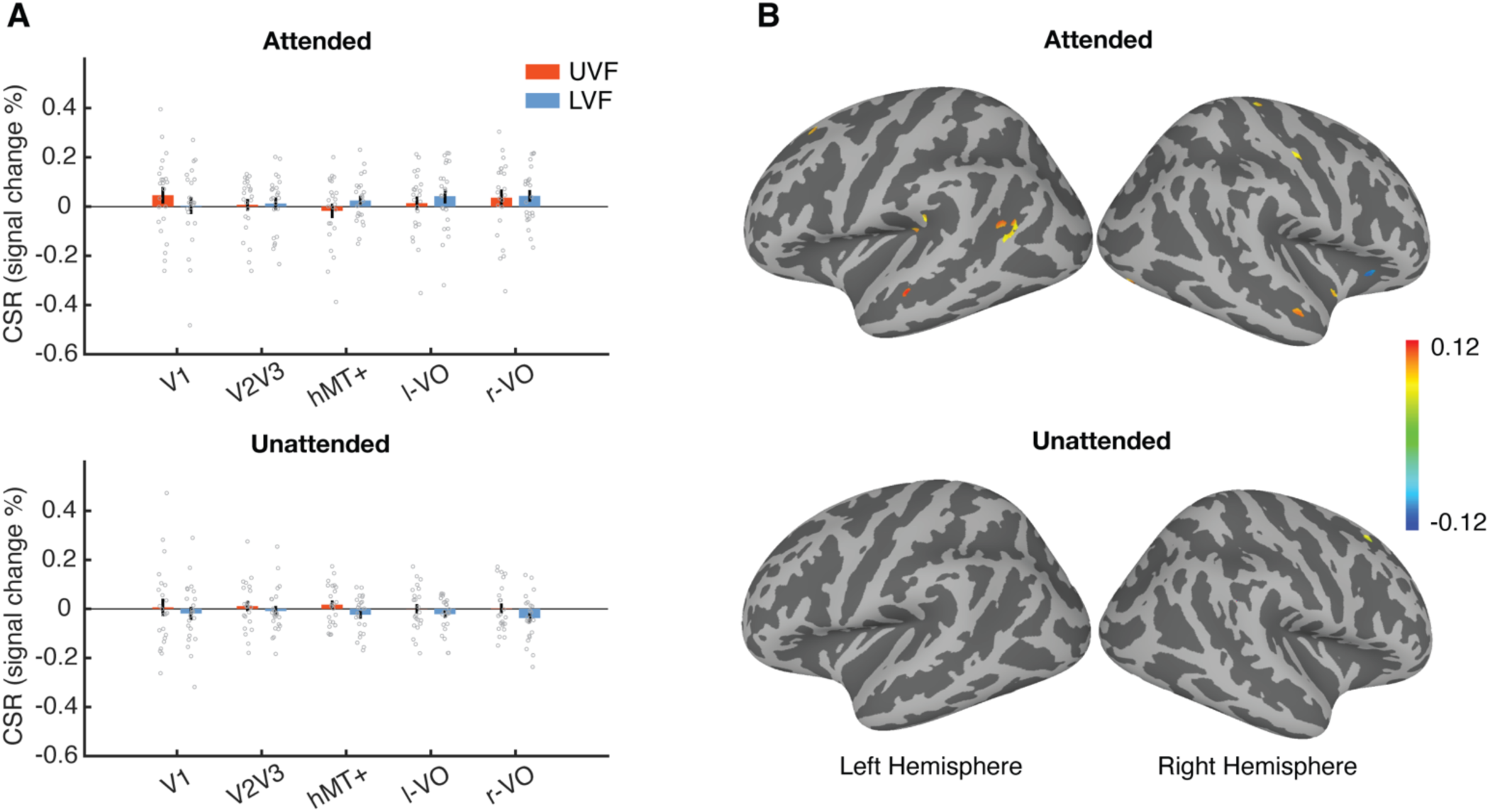
Collision sensitive response (CSR) in cortical areas. (A) CSR, defined as response difference between the hit and miss conditions, in early visual areas. (B) CSR in other cortical regions. The color bar indicates CSR in percent signal change. Maps were presented with a threshold of uncorrected p<0.001. No clusters survived following FWE correction.

The results were comparable when the responses in the upper and lower visual fields were combined or when they were analyzed separately (data not shown). hMT+, human middle temporal complex; l/r-VO, left/right ventral occipitotemporal cortex. Other conventions as in Fig. 2.

## 3. Divergent subcortical pathways exhibit collision sensitivity depending on the state of attention and consciousness

To reveal the subcortical pathways underlying collision detection, we used correlation and path analysis of collision sensitive response (CSR), and beta series connectivity analysis (Rissman et al., 2004; Cisler et al., 2014) to investigate the information flow in subcortical regions. A significant correlation was found between CSRs in the superficial SC and PBGN in the ATT-UVF condition (r = 0.446, p = 0.033, Fig. 5A), but not in the other conditions (all r < 0.2). Path analysis was performed using CSR correlations with structural equation modeling (SEM) (Zhuang et al., 2005; Jun et al., 2020). In the effective connectivity model (Fig. 5B), connections among the superficial SC, PBGN, pulvinar, VTA, and CMA were defined by the anatomical projections in primates (May, 2006; Rafal et al., 2015; Zubair et al., 2021) and established neural circuits in rodents (Shang et al., 2015; Wei et al., 2015; Zhou et al., 2019). For each attention × visual field condition, the covariance matrix among ROIs was used to fit the SEM to calculate the connectivity strength (Fig. 5C, Fig. S6). In the ATT-UVF condition, significant positive connectivity was observed for the SC-PBGN (connectivity strength (β) = 0.45, p = 0.017) and SC-VTA (β = 0.41, p = 0.029) pathways (see Fig. S6 for other conditions). Moreover, the SC-PBGN connection was significantly stronger in the attended than in the unattended condition (p < 0.001, UVF), suggesting that top-down attention significantly enhanced collision sensitivity in the SC-PBGN pathway to looming stimuli in the upper visual field.

**Fig. 5.**
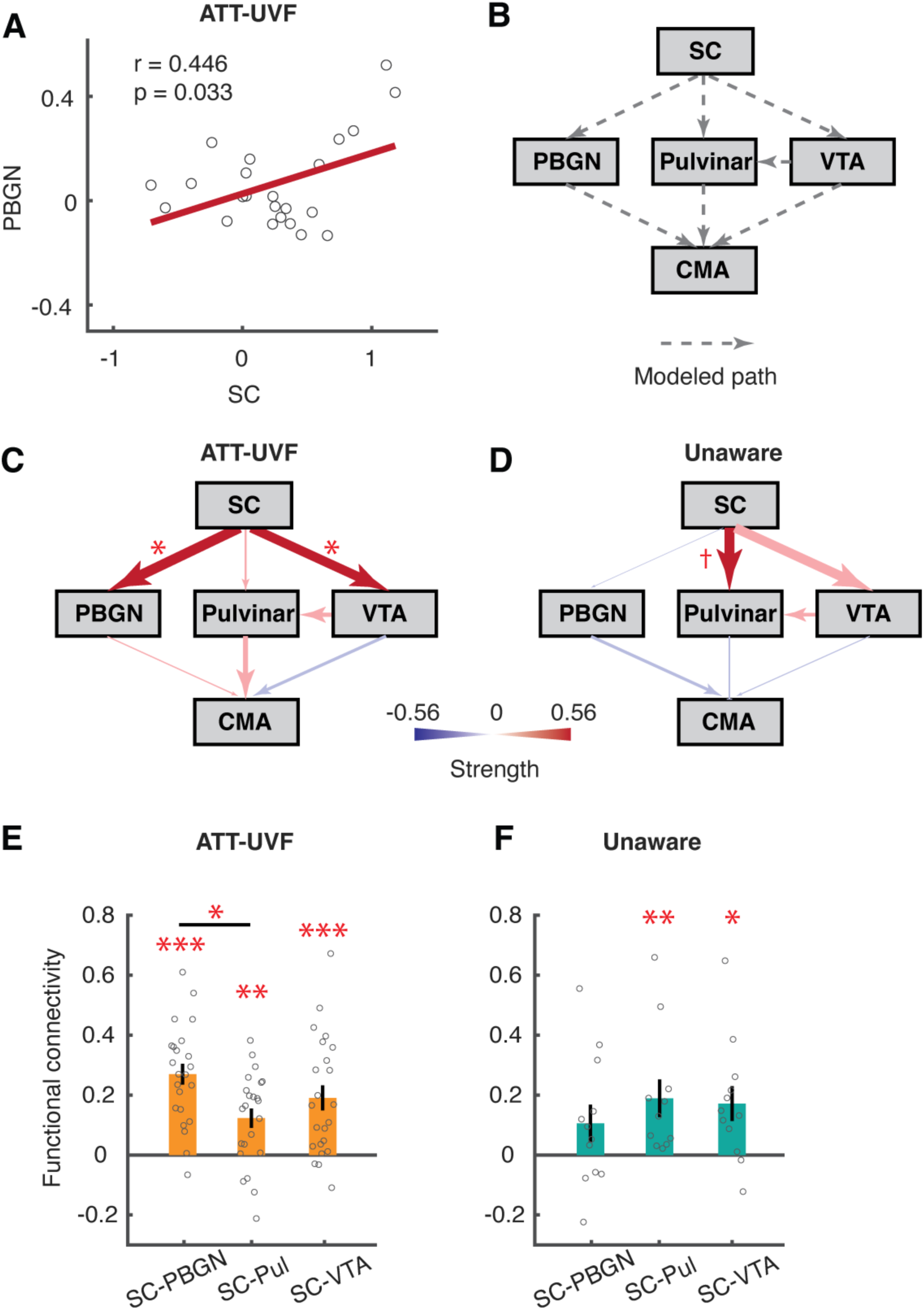
Correlation and connectivity analyses in the subcortical regions. (A) Scatterplot of collision sensitive responses (CSR) in the PBGN and the superficial SC in the ATT-UVF condition. Red line indicates linear fit of individual data (gray circles). (B) The structural equation model (SEM) for the path analysis. Gray dashed arrows indicate connections based on known anatomical projections. (C) SEM results in the ATT-UVF condition in healthy participants. (D) SEM results for looming stimuli presented in the blind visual field of hemianopic patients. Dark and light colors depict significant and non-significant connections, respectively. (E, F) Beta series functional connectivity in the ATT-UVF condition for healthy participants and unaware condition for hemianopic patients. The connectivity was assessed using beta series for the hit trajectories. *** p<0.001; ** p<0.01; * p<0.05; † p<0.1.

To further investigate the subcortical pathways for collision detection without awareness to looming stimuli, we re-analyzed the fMRI data from hemianopic patients with unilateral lesions of the geniculostriate pathway in our previous study (Guo et al., 2024). Visual stimuli were similar to the hit-eye and near-miss conditions in the current study except that they were presented on a 2-D screen. For collision sensitive responses to looming stimuli presented in the blind visual field, we observed a trend of positive correlation between SC and Pulvinar (r = 0.539, p = 0.087), and between SC and VTA (r = 0.455, p = 0.138). Path analysis further revealed a marginal effect for SC-Pulvinar (β = 0.44, p = 0.095), and SC-VTA (β = 0.42, p=0.102) connections. In contrast, the SC-PBGN connection was either weak or absent (β = -0.03, p=0.88) (Fig. 5D). Together with the findings from healthy participants (Fig. 5C), these results suggest that divergent subcortical pathways are differently engaged in collision detection depending on the state of attention and consciousness.

The beta series connectivity for hit responses to UVF stimuli exhibited qualitatively similar pattern of results. For the attended condition in healthy participants (Fig. 5E), the SC-PBGN pathway showed the strongest connectivity strength (F(2, 44) = 4.51, p = 0.017; SC-PBGN > SC-Pul: p = 0.014). For stimuli presented in the blind visual field of hemianopic patients (unaware condition, Fig. 5F), a significant connectivity was observed for the SC-Pulvinar (p = 0.004) and SC-VTA (p = 0.015) pathways. While the SC-PBGN connectivity was not significant and weaker than that in the attended condition (uncorrected p = 0.018).

Taken together, these findings suggest that SC-Pulvinar and SC-VTA pathways may support collision detection without attention and awareness, whereas the SC-PBGN pathway mediates accurate estimation of collision trajectory under top-down attentional control.

## Discussion

A number of subcortical nuclei, including the superficial SC, PBGN, and centromedial amygdala, exhibited sensitivity to collision trajectories under top-down attention, but not when attention was diverted away from the looming stimuli. Collision sensitivity was strongest for hit-nasion trajectories presented in the upper visual field. Correlation and path analysis suggest that the SC-PBGN pathway mediates collision detection under top-down attention. In contrast, the SC-pulvinar and SC-VTA pathways may support collision detection without attention and awareness to stimuli presented in the blind visual field of hemianopic patients.

Neural circuits formed by parvalbumin-positive (PV+) excitatory neurons in the superficial SC, along with their downstream targets in the PBGN and amygdala, have been shown to mediate defensive responses to looming stimuli in rodents (Shang et al., 2015; Shang et al., 2018). The current study provides further evidence that the SCs-PBGN pathway also detects collision trajectories in humans, but only when participants are paying attention to the looming stimuli. Moreover, collision-sensitive responses were stronger for the hit-nasion than the hit-eye trajectory, suggesting that binocular information was utilized by the subcortical pathway for accurate estimation of collision trajectories. In support of this explanation, neurons in the primate SC receive binocular retinal inputs (Dilbeck et al., 2022), and some tuned to opposite motion directions from the two eyes (Tailby et al., 2012), consistent with the retinal motion produced by the hit-nasion trajectory. A binocular region in the rodent SC connected with the PBGN also facilitates detecting looming stimuli in the UVF (Deichler et al., 2020). Thus, it is possible that top-down attention may selectively enhance the sensitivity of binocular neurons in the SC-PBGN pathway to improve the accuracy of collision detection.

Our previous study showed stronger SC responses to near-miss trajectories in the UVF than those in the LVF (Guo et al., 2024), whereas no significant difference was observed for collision trajectories, likely due to response saturation at short TTC. Using looming stimuli at a longer TTC, the current study revealed a clear UVF advantage in subcortical responses to collision compared to near-miss trajectories, aligned with behavioral findings of higher collision sensitivity in the UVF than in the LVF (Guo et al., 2024). Consistent with our human results, the tectofugal pathways in rodents also mediate defensive responses particularly to looming stimuli in the UVF (Wei et al., 2015; Zhou et al., 2019; Deichler et al., 2020; Isa et al., 2020). Moreover, the SC in non-human primates also overrepresents the UVF with sharper, stronger and faster neural responses (Hafed and Chen, 2016). Together, these findings suggest that tectal pathways are phylogenetically conserved and ecologically adapted for detecting threats from the UVF, such as aerial predators.

In contrast to the attention-gated function of the SC-PGBN pathway, path analysis suggests that SC-Pulvinar and SC-VTA pathways may be involved in automatic detection of collision trajectories without attention and awareness to the looming stimuli in blind visual field of hemianopic patients (Fig. 5D). This finding is consistent with the collision-sensitive activations previously observed in these regions (Guo et al., 2024). These subcortical pathways may serve a “low road” shortcut for processing threatening, emotional and social information in exteroceptive input (Pessoa and Adolphs, 2010; Kragel et al., 2021). In addition to collision detection without attention and awareness, the SC–VTA pathway may also contribute to collision detection under attention (Fig. 5C). The absence of collision-sensitive responses in the VTA may reflect its high sensitivity to all looming stimuli, regardless of trajectory (Fig. 3B). In rodents, divergent tectofugal and thalamic pathways have been shown to mediate different types of defensive behaviors, such as escape and freezing (Salay et al., 2018; Shang et al., 2018). Our data suggest that in humans, subcortical pathways are differently engaged in collision detection depending on the state of attention and consciousness: the SC-PBGN pathway, under top-down attention, can integrate binocular information for precise estimation of collision trajectories; whereas the SC-pulvinar and SC-VTA pathway support collision detection without attention and awareness.

## Methods

### Participants

A total of 23 healthy young adults (9 males, 21 naive participants) participated in the experiment. This sample size provides 80% power (Type I error < 0.05) for detecting an effect size (Cohen’s d) greater than 0.611 for the key comparison between hit-nasion and near-miss trajectories in the current study. The age of the participants ranged from 20 to 40 years old, with normal or corrected-to-normal vision. All participants provided written informed consent before the experiments. The Institutional Review Board of the Institute of Biophysics approved the experimental protocols (Ethic No. 2012-IBP-011). The data of 12 hemianopic patients with unilateral lesions of the geniculostriate pathways from our previous study (Guo et al., 2024) were re-analyzed for the path analysis. Patients lost their conscious vision in one-half or a quadrant of the visual field (relative sensitivity less than -20 dB and p < 0.5% compared with normal population in both eyes), while they had normal or corrected-to-normal vision outside the blind visual field. For these patients, the clinical characteristics, stimuli and procedure, and MRI data acquisition and preprocessing, were described in our previous study.

### Apparatus, stimuli and procedure

Visual stimuli were generated by MATLAB with Psychtoolbox 3.0 (Brainard, 1997; Pelli, 1997). 3D spheres, rendered from slightly different perspectives, were presented to the two eyes with a MRI compatible polarized LCD display (3D BOLD screen 24, Cambridge Research Systems) and polarized glasses. As shown in Figure S1, the 3D sphere appeared at 11.3 m and vanished at 3.3 m from the observer, moving at a speed of 24 m/s. The time-to-collision at the last frame was 138 ms. The trajectory started in one of the four quadrants of visual field, with a 0.65-m horizontal offset and a 0.38-m vertical offset. It would either hit the nasion (hit nasion), hit the ipsilateral eye (hit eye, 3 cm from nasion), or slightly miss the head (near miss) of observers. The near-miss trajectory was adjusted for each individual to achieve a 50% hit response in a training session outside the scanner. To match their retinotopic locations, the on-screen projections of looming stimuli were aligned to the center of mass. The on-screen image expanded from 0.3 to 1 degree of visual angle in 330 ms, with a mean eccentricity of 3.8 degrees. Dark stimuli (mean luminance = 2.9 cd/m^2^) were presented on a bright background (5.8 cd/m^2^) in 11 participants, while bright stimuli (mean luminance = 5 cd/m^2^) were presented on a black background (∼0 cd/m^2^) in 12 participants.

A long event-related design was adopted for the fMRI experiment, with inter-stimulus intervals (ISIs) of 10, 12, or 14 seconds. A total of 48 trials were collected for each trajectory × attention condition, with 12 trials in each quadrant of visual field. In the attended condition, participants were instructed to maintain fixation while making judgement whether the incoming object was on a collision course with their head. In the unattended condition, they performed a rapid RSVP task at fixation. In which, they detected letter targets (e.g., white ‘T’ and blue ‘O’) among letter distractors. In the retinotopic localizer, a checkerboard disc (1.9 degrees in diameter) counter-phase flickering at 7.5 Hz was presented in 500-ms on/off for 8 seconds at one location, followed by 8-second fixation period. The stimulus cycled through the four quadrant locations corresponding to the looming stimuli in separate blocks. A total of 16 blocks of data were collected for each location. Participants were instructed to maintain fixation while passively viewing the stimuli. Two sessions were collected for the looming experiment, and another session for the retinotopic localizer (4 runs, each for 256 s). In 11 participants, fMRI data for the attended (10 runs, each for 242 s) and unattended (12 runs, each for 242 s) conditions were collected in separate sessions. The RSVP stimuli were presented only in the unattended condition. For the remaining 12 participants, attended (8 runs, each for 288 s) and unattended (8 runs, each for 288 s) conditions were scanned in odd and even runs within each session, with the RSVP stimuli presented in both conditions.

For hemianopic patients, similar stimuli were presented on a 2D screen. In different trials, a sphere moved along hit (hit-eye), near-miss, or receding (backward-moving) trajectories. The TTC of the last frame of the looming stimuli was 115 ms (2.75 m from the observer). Stimuli were presented in either the blind or normal visual field of patients. During fMRI scans, patients performed a central fixation task irrelevant to the moving sphere. Subcortical fMRI responses to stimuli in the blind visual field in the hit and near-miss conditions were obtained for path analysis in the current study. Refer to Guo et al. (2024) for more details.

### MRI data acquisition and preprocessing

MRI data was acquired using a 3T MRI scanner (Siemens Prisma Fit, Erlangen, Germany) and a 20-channel head coil (Nova Medical, Cambridge, MA, USA) at Beijing MRI center for Brain Research. Functional images were acquired with a gradient recalled echo planar imaging (GE-EPI) sequence (2-mm isotropic voxels, 54 axial slices, 192-mm FOV, 96 × 96 matrix, TR/TE = 2000/31.4 ms, flip angle = 80°, multiband factor = 2, no parallel imaging). High-resolution anatomic images were obtained with a T1w MPRAGE sequence (1-mm isotropic voxels, 192 sagittal slices, 256 × 256 matrix, TR/TE = 2600/3.02 ms, flip angle = 8°). Pulse and respiration data were recorded for the first 11 participants.

Imaging data were analyzed using AFNI (Cox, 1996) and Freesurfer (Dale et al., 1999; Fischl et al., 1999). The pre-processing steps included: RETROICOR physiological noise removal (Jo et al., 2010); slice timing correction; motion correction and co-registration of functional images to T1w anatomical volumes; functional data resampled (cubic interpolation) at 1-mm isotropic resolution; scaled to percent signal change. In volume-based analysis of subcortical data, nonlinear transformation to the standard space was focused on subcortical regions using a weight mask (AFNI 3dQWarp). For the retinotopic localizer, 2-mm FWHM spatial smoothing was performed within subcortical ROIs (AFNI 3dBlurinMask). In surface-based analysis of cortical data, functional volumes following motion correction and co-registration were mapped onto the cortical surface, and smoothed on the surface with a 6-mm FWHM gaussian kernel. Surface data were spatially normalized based on a standard-mesh method (Argall et al., 2006). After pre-processing, a general linear model (GLM) with a canonical HRF (BLOCK4(0.5,1) in AFNI) was used to estimate the response (beta value) to each stimulus condition. Baseline regressors included zero-, first-and second-order polynomials. Noise regressors included 6 rigid motion parameters, and a timecourse from the anterior cerebellum to reduce physiological noise in subcortical regions, especially the SC (Wall et al., 2009). For accurate estimation of collision-sensitive response in the SC, a faster HRF with an early time to peak (4s), early fall time and moderate undershoot fraction (WAV(0,1,3,4.2,0.5,2.8) in AFNI) was used in the GLM (Wall et al., 2009; DeSimone et al., 2015). Event-related average of response timecourse in the SC was calculated using the detrended time series (AFNI 3dTproject), after removing baseline and noise regressors. The signal intensity at stimulus onset and 2s before were used as the baseline.

### ROI definition

The whole anatomical ROIs of the SC were manually drawn on T1w anatomical images. The depth of each voxel was defined as the minimum distance to the surface of the SC (Fig. 2A). Voxels of 0 to 1.5 mm depth were defined as the superficial layers, and those of 1.5 to 3 mm depth as the intermediate layers. Retinotopic ROIs of the SC were defined as the most significant 15 μL of voxels (p<0.01 for all voxels) with contralateral-ipsilateral activations, separately for the UVF and LVF. Different volumes of the selected ROIs led to qualitatively similar conclusions for the subsequent analysis (Fig. S2). LGN and pulvinar ROIs were define by a combination of anatomical constraint and significant localizer activation (voxel p < 0.02, the threshold was selected to ensure the cluster size > 10 µL) (Arcaro et al., 2015; DeSimone et al., 2015). The whole anatomical ROIs were used for the PBGN and VTA. PBGNs are located bilaterally in the midbrain, below the inferior colliculus and above the substantia nigra (Pzxinos, 2012; Mai, 2016). VTA ROIs were manually defined from the midbrain on T1w anatomical images refer to a previous study (Ballard et al., 2011). For retinotopic regions including the SC, LGN and pulvinar, their responses to contralateral looming stimuli were analyzed. While for PBGN and VTA, responses to both hemifields were averaged.

The amygdala complex was divided into a basolateral (BLA) and a centromedial (CMA) part based on the probabilistic cytoarchitectonic atlas derived from human post-mortem studies (Amunts et al., 2005; Eickhoff et al., 2005). Given that the neural circuits from the SC to amygdala have been shown mediating defensive behaviors to looming stimuli (Pegna et al., 2005; Rafal et al., 2015; Shang et al., 2015; Wei et al., 2015; Koller et al., 2019), BLA and CMA ROIs were defined as voxels showing significant functional connectivity with the SC using a beta-series method (refer to functional connectivity section) (Rissman et al., 2004). In which, the response for each trial was estimated with a separate regressor in a GLM, generating a series of beta values. Beta values were Z-scored within each condition. A linear regression was conducted with the beta series from the target (SC) and source (BLA/CMA) ROIs, and the regression coefficient was obtained as the strength of functional connectivity. A leave-one-subject-out (LOSO) procedure was used to define the ROIs for each participant. BLA and CMA ROIs were defined as the largest significant cluster in the nucleus from the group-level connectivity map from all other participants (voxel p < 0.02, cluster size > 10 μL).

Frontoparietal ROIs were defined on the standard surface (std.141 in AFNI) using HCP-MMP1 atlas (Glasser et al., 2016). ROIs for visual cortices were defined by significant activations in the retinotopic localizer (p<0.05 after FDR correction). V1-V3 ROIs were defined by localizer vs. baseline in each quadrant, respectively. hMT+ were defined by contralateral vs. ipsilateral. VO ROIs were defined by the average of all localizer stimuli vs. baseline.

For hemianopic patients, in the absence of a retinotopic localizer, a leave-one-run-out (LORO) procedure was used to define the SC and pulvinar ROIs for the path analysis. Within the anatomical ROIs, voxels most responsive to collisions (16-17.28 μL, depending on fMRI resolution, hit-miss p < 0.05 for all voxels) in N-1 runs were used to calculate the ROI-averaged response for the left-out run. The procedure was repeated for each run and the results were averaged across all runs. Since collision-sensitive activation was mainly observed in the ventromedial pulvinar, this sub-region was used as the anatomical ROI for the pulvinar in the LORO analysis. ROI definitions for the PBGN, VTA and CMA were the same as those used for healthy participants.

### Path analysis

Path analysis with SEM methods was performed to estimate the effective connectivity. The CSRs of all participants were input into a fixed model of connections based on prior anatomical evidence. To avoid the influence of mean response levels on the estimated connection strength, CSRs were Z-scored for each ROI and condition. Then the path coefficients were fitted by maximum likelihood estimate in the SEM with lavaan software 0.6-17 (Rosseel, 2012; Merkle and Rosseel, 2018). The path coefficients were compared between different conditions using two-sample Z tests:

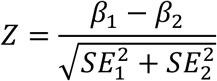

in which *β* and SE denote the strength and standard error of the estimated connection.

### Functional connectivity

A beta-series method was adopted to measure the functional connectivity between subcortical regions (Rissman et al., 2004). The response for each trial was estimated with a separate regressor in a GLM model of the ROI-averaged BOLD time course, generating a series of beta values. The beta values for each condition were Z-scored to avoid the influence of mean activation. Then the correlation coefficient between two beta-series from different ROIs was obtained as the functional connectivity.

### Statistics

We performed LME analysis for the ROI-averaged responses separately for the attended and unattended conditions. Trajectory (hit nasion, hit eye, or near miss), visual field (UVF or LVF), and stimulus luminance (bright or dark; between-subject factor) were included as fixed effects, with participant as a random grouping factor. Collision sensitivity was first assessed as an interaction effect between trajectory and visual field or a main effect of trajectory. Then, the trajectory effects in the UVF and LVF were tested in separate LME models. Finally, post hoc paired t-tests with Holm corrections were performed between different trajectory conditions if a significant effect of trajectory was observed.

Whole-brain analysis was performed on the cortical surface to detect potential collision sensitivity without making specific prior assumptions on the ROIs. At a threshold of p < 0.001, group-level activations were corrected for family-wise error (p < 0.05) using a cluster size of 140 mm^2^. The cluster-size threshold was obtained using a simulation procedure with self-generated all-noise data on the same surface model and blurring level (AFNI slow_surf_clustsim.py, iteration = 10000000).

Functional connectivity was first obtained for each individual. Then the significance of connectivity was examined with non-parametric exact permutation test considering the skewed distribution of connectivity data (same as correlation coefficient). The p values were further corrected for multiple comparison (6 comparisons) problems with Holm method. For each subject group (healthy subjects in ATT-UVF conditions, patients in unaware conditions), we also compared the connectivity between different pathways using LME models with subject as a random effect. Fisher transformation was applied to the connectivity data before the LME analysis. Following paired test were performed using exact permutation test with Holm correction if a significant effect of pathways were observed. Finaly, we compared the connectivity for the same pathways between the two subject groups using permutation test (permutation=10000000) for independent samples with Holm correction (3 comparisons).

## Data and code availability

The raw MRI data from this study will be made publicly available at Openneuro.org upon publication. The custom codes for ROI analysis are deposited at Zenodo.org and publicly available (https://zenodo.org/records/17225019).

## Author Contributions

J.Z. and P.Z. designed research; J.Z and Y.W. performed research; J.Z., Y.W. and P.Z. analyzed data; J.Z. and P.Z. wrote the paper.

## Acknowledgments

This study was supported by STI2030-Major Projects released by the Ministry of Science and Technology of China (2022ZD0211900 and 2021ZD0204200, https://service.most.gov.cn/index/), National Natural Science Foundation of China (31871107 and 31930053, https://www.nsfc.gov.cn/english/site_1/index.html) to P.Z., Provincial Natural Science Foundation of Hunan (2024JJ9001, https://kjt.hunan.gov.cn/zxgz/zkjj/), Hunan Xiangjiang Philanthropy Foundation (KY24016, http://hnxjgy.cn) and Clinic Research Foundation of Aier Eye Hospital Group (AIM2301D04, http://kjgl.aierchina.com:8088/home) to J.Z.. The funders had no role in study design, data collection and analysis, decision to publish, or preparation of the manuscript.

## SI figures

**Fig. S1.**
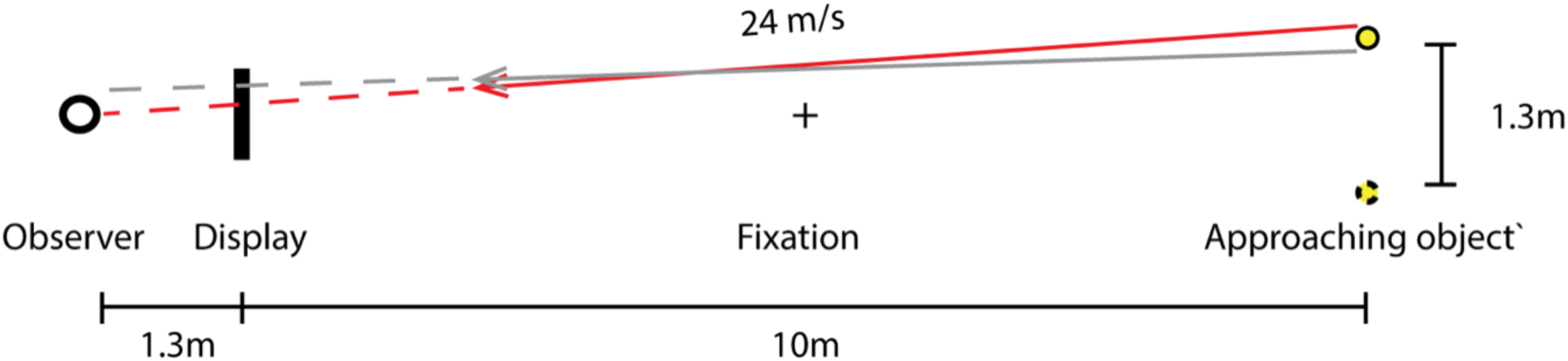
Top view of the stimulus trajectories. This schematic is a scaled-down version of the scenes simulated in the experiment. Only hit-nasion (red) and near-miss (gray) trajectories wereplotted. The point of collision and passage were indicated by the extended dashed lines.

**Fig. S2.**
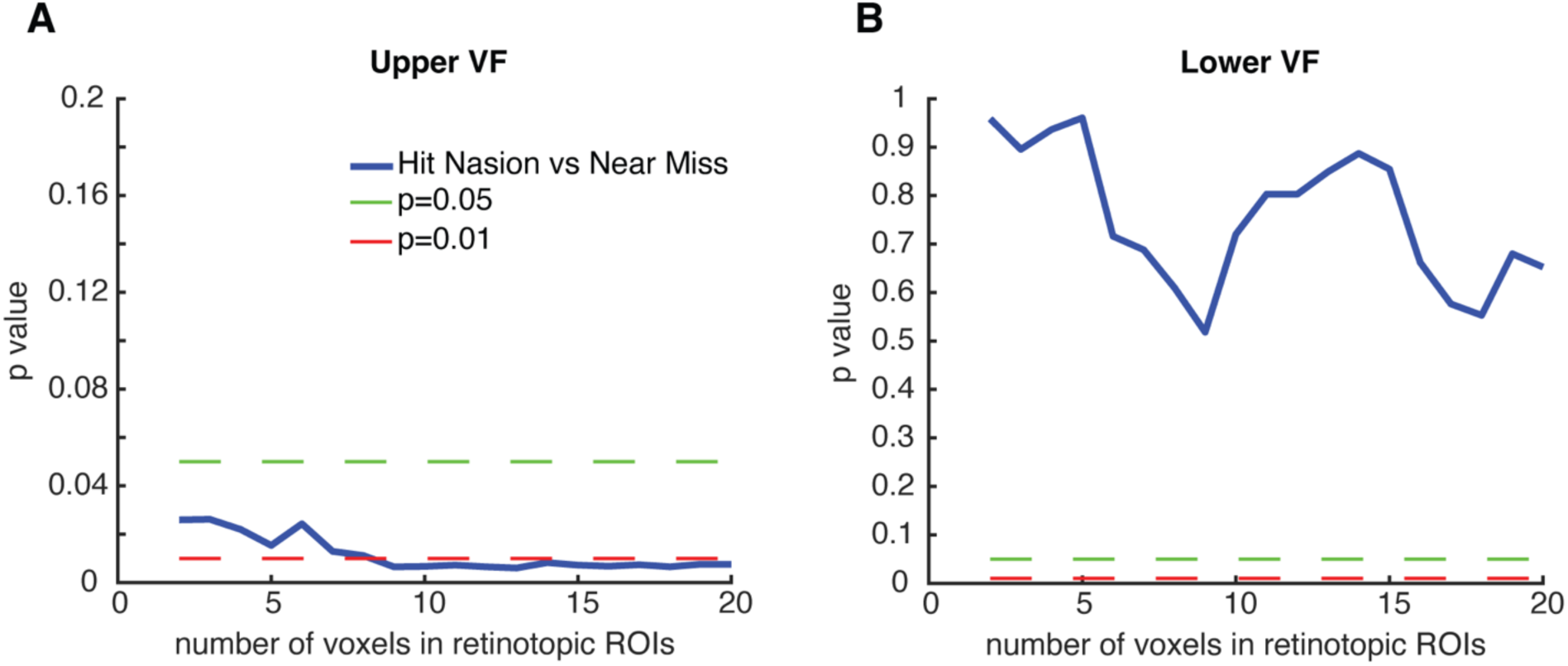
Statistical significance (p-values) of collision sensitivity as a function of voxel number in the retinotopic ROIs of the superficial SC. The number of voxels in the retinotopic ROIs was controlled by the threshold of contralateral vs. ipsilateral activation in the localizer. Collision sensitivity (hit nasion vs. near miss in the attended conditions) remained significant in the upper (A) visual field, and insignificant in the lower (B) visual field at different ROI-selection thresholds.

**Fig. S3.**
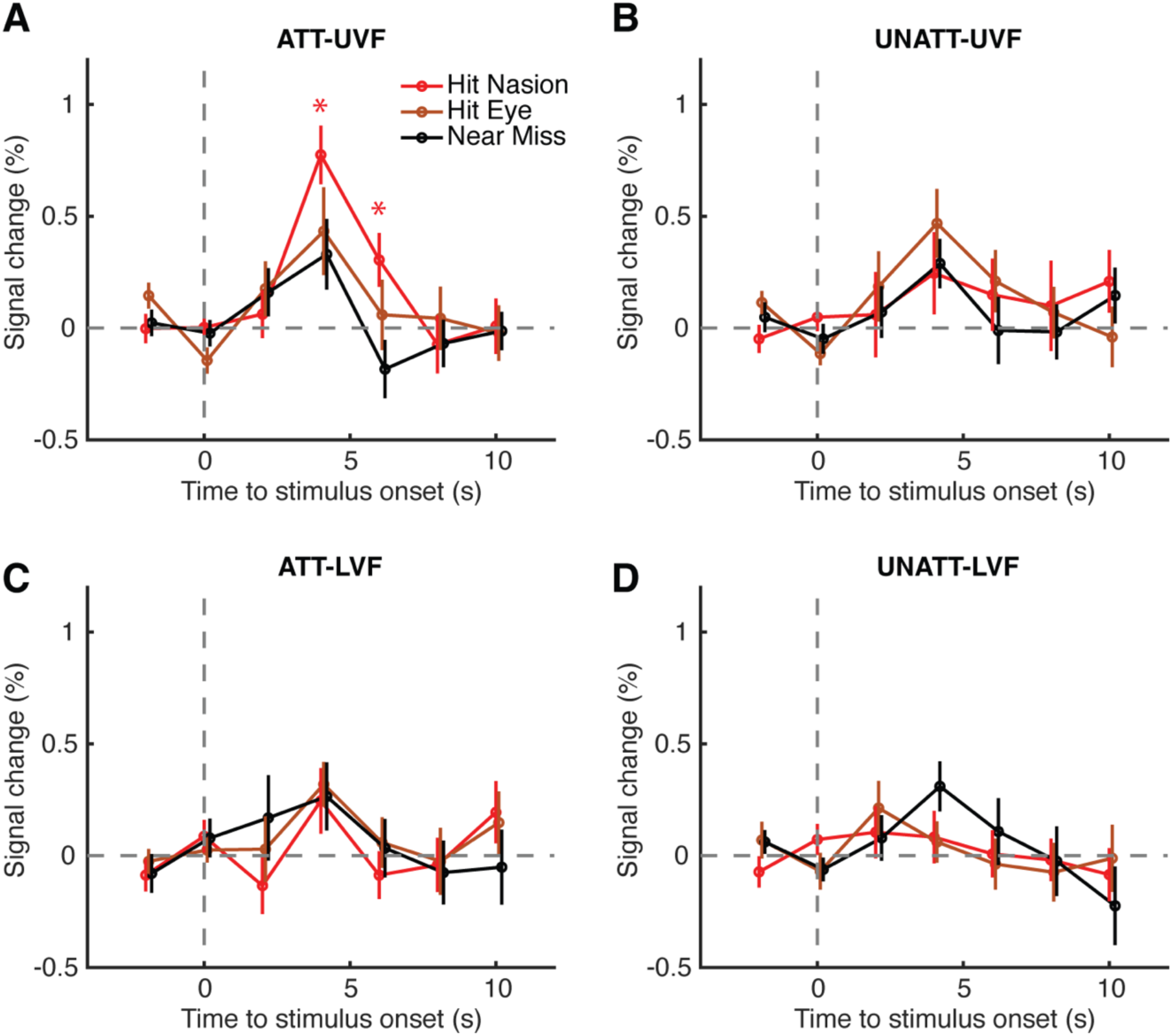
Event-related response time series in the superficial SC. * denotes p<0.05 for hit-nasion vs. near-miss conditions. Error bars indicate S.E.M. Abbreviations: ATT, attended; UNATT, unattended; UVF, upper visual filed; LVF, lower visual field.

**Fig. S4.**
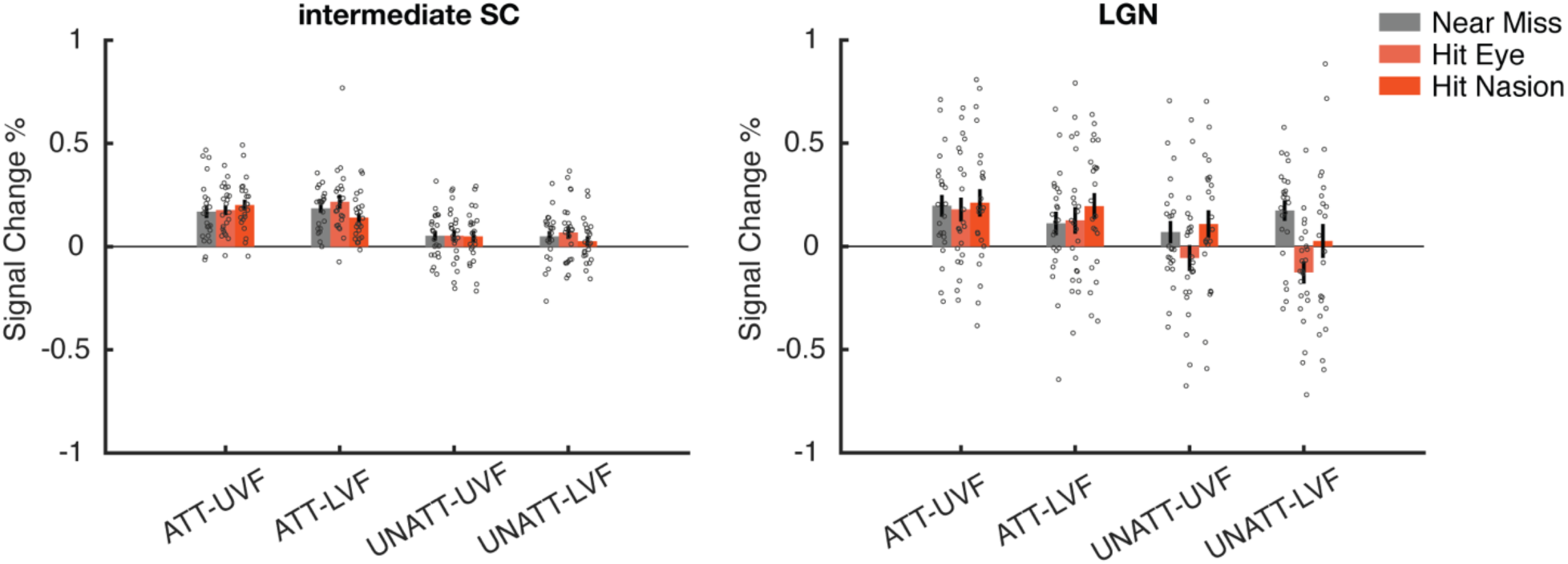
Looming responses in the intermediate layers of the SC and the LGN. Dots denote the data of individual participants. Error bars indicate S.E.M. ATT, attended; UNATT, unattended; UVF, upper visual filed; LVF, lower visual field.

**Fig. S5.**
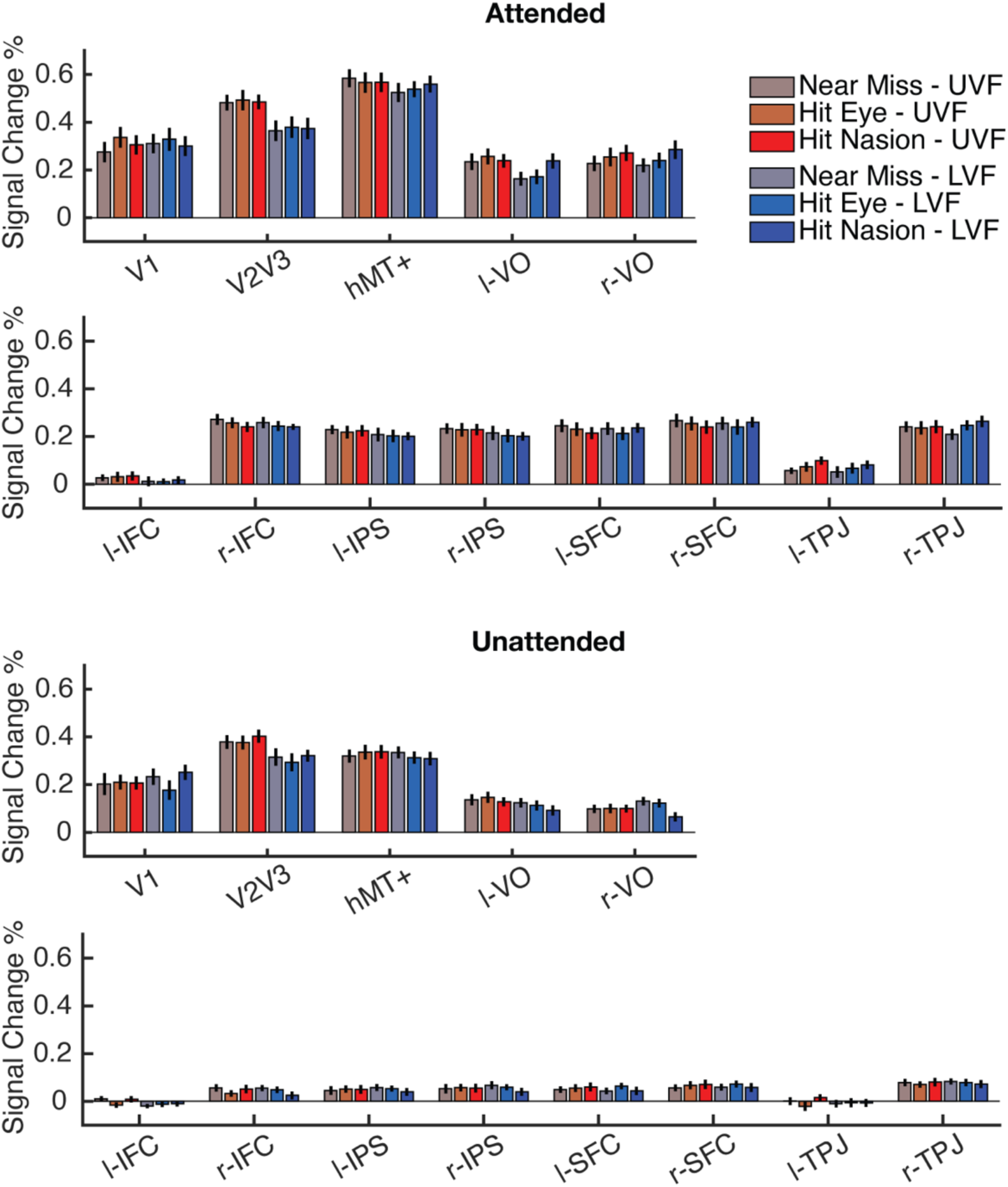
Looming responses in the visual cortices and frontoparietal attention networks. UVF, upper visual filed; LVF, lower visual field. hMT+, human middle temporal complex; VO, ventral occipital cortex; IFC, inferior frontal cortex; IPS, intraparietal sulcus; SFC, superior frontal cortex; TPJ, temporoparietal junction. l/r-, left/right hemisphere.

**Fig. S6.**
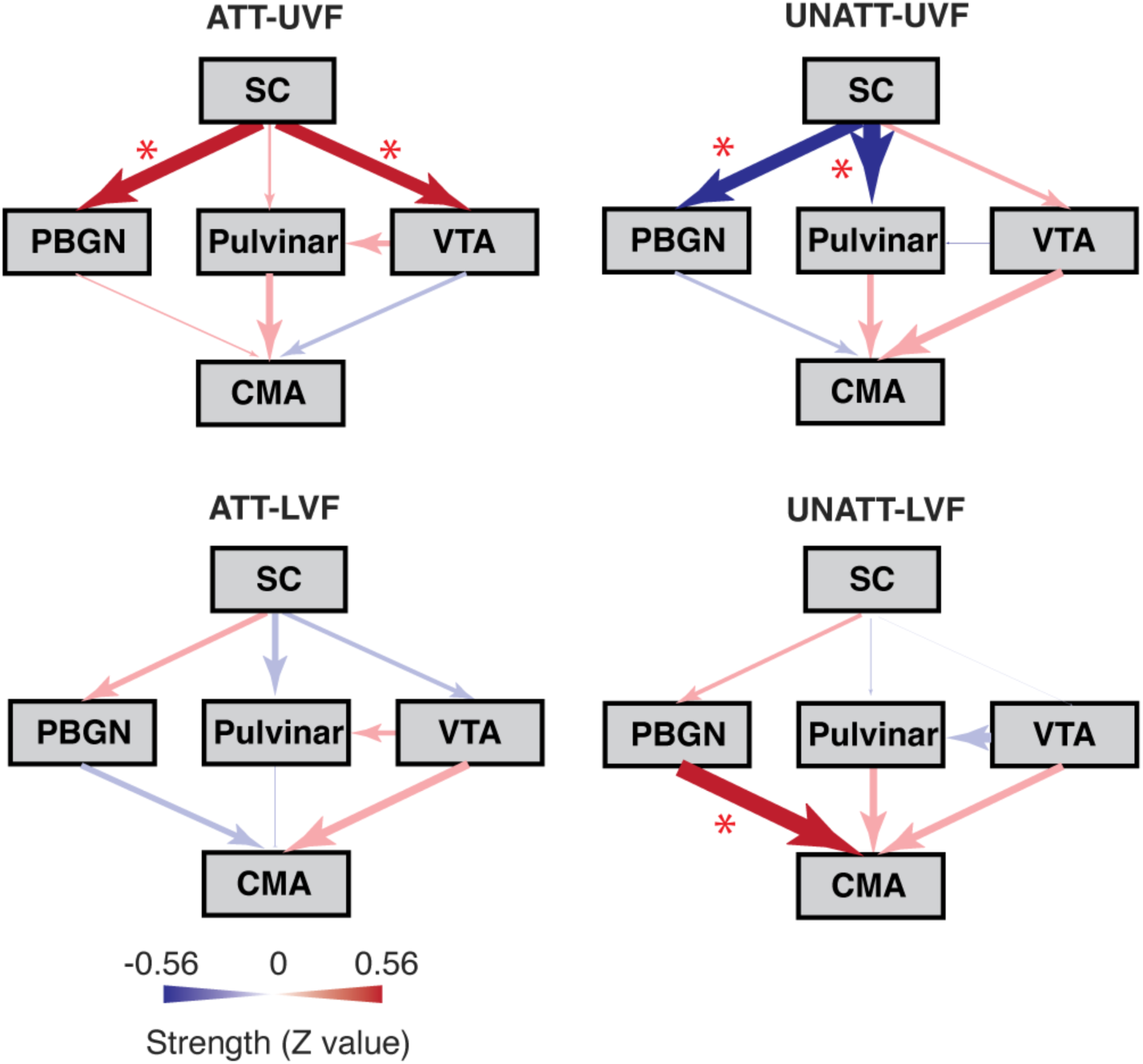
Path analysis results in all stimulus conditions. Dark and light colors depict significant and non-significant connections, respectively. ATT, attended; UNATT, unattended; UVF, upper visual filed; LVF, lower visual field. * p<0.05.

## References

Amunts K, Kedo O, Kindler M, Pieperhoff P, Mohlberg H, Shah NJ, Habel U, Schneider F, Zilles K (2005) Cytoarchitectonic mapping of the human amygdala, hippocampal region and entorhinal cortex: intersubject variability and probability maps. Anat Embryol (Berl) 210:343–352.

Arcaro MJ, Pinsk MA, Kastner S (2015) The Anatomical and Functional Organization of the Human Visual Pulvinar. J Neurosci 35:9848–9871.

Argall BD, Saad ZS, Beauchamp MS (2006) Simplified intersubject averaging on the cortical surface using SUMA. Hum Brain Mapp 27:14–27.

Ballard IC, Murty VP, Carter RM, MacInnes JJ, Huettel SA, Adcock RA (2011) Dorsolateral prefrontal cortex drives mesolimbic dopaminergic regions to initiate motivated behavior. J Neurosci 31:10340–10346.

Billington J, Wilkie RM, Field DT, Wann JP (2011) Neural processing of imminent collision in humans. Proc Biol Sci 278:1476–1481.

Brainard DH (1997) The Psychophysics Toolbox. Spat Vis 10:433–436.

Chen L, Yuan X, Xu Q, Wang Y, Jiang Y (2016) Subliminal Impending Collision Increases Perceived Object Size and Enhances Pupillary Light Reflex. Front Psychol 7:1897.

Cisler JM, Bush K, Steele JS (2014) A comparison of statistical methods for detecting context-modulated functional connectivity in fMRI. Neuroimage 84:1042–1052.

Cox RW (1996) AFNI: software for analysis and visualization of functional magnetic resonance neuroimages. Comput Biomed Res 29:162–173.

Cynader M, Berman N (1972) Receptive-field organization of monkey superior colliculus. J Neurophysiol 35:187–201.

Dale AM, Fischl B, Sereno MI (1999) Cortical surface-based analysis. I. Segmentation and surface reconstruction. Neuroimage 9:179–194.

Deichler A, Carrasco D, Lopez-Jury L, Vega-Zuniga T, Márquez N, Mpodozis J, Marín GJ (2020) A specialized reciprocal connectivity suggests a link between the mechanisms by which the superior colliculus and parabigeminal nucleus produce defensive behaviors in rodents. Sci Rep 10:16220.

DeSimone K, Viviano JD, Schneider KA (2015) Population Receptive Field Estimation Reveals New Retinotopic Maps in Human Subcortex. J Neurosci 35:9836–9847.

Dilbeck MD, Spahr ZR, Nanjappa R, Economides JR, Horton JC (2022) Columnar and Laminar Segregation of Retinal Input to the Primate Superior Colliculus Revealed by Anterograde Tracer Injection Into Each Eye. Invest Ophthalmol Vis Sci 63:9.

Ding SL et al. (2016) Comprehensive cellular-resolution atlas of the adult human brain. J Comp Neurol 524:3127–3481.

Duke PA, Rushton SK (2012) How we perceive the trajectory of an approaching object. J Vis 12:9.

Eickhoff SB, Stephan KE, Mohlberg H, Grefkes C, Fink GR, Amunts K, Zilles K (2005) A new SPM toolbox for combining probabilistic cytoarchitectonic maps and functional imaging data. Neuroimage 25:1325–1335.

Fischl B, Sereno MI, Dale AM (1999) Cortical surface-based analysis. II: Inflation, flattening, and a surface-based coordinate system. Neuroimage 9:195–207.

Glasser MF, Coalson TS, Robinson EC, Hacker CD, Harwell J, Yacoub E, Ugurbil K, Andersson J, Beckmann CF, Jenkinson M, Smith SM, Van Essen DC (2016) A multi-modal parcellation of human cerebral cortex. Nature 536:171–178.

Guo F, Zou J, Wang Y, Fang B, Zhou H, Wang D, He S, Zhang P (2024) Human subcortical pathways automatically detect collision trajectory without attention and awareness. PLoS Biol 22:e3002375.

Hafed ZM, Chen CY (2016) Sharper, Stronger, Faster Upper Visual Field Representation in Primate Superior Colliculus. Curr Biol 26:1647–1658.

Hubel DH, LeVay S, Wiesel TN (1975) Mode of termination of retinotectal fibers in macaque monkey: an autoradiographic study. Brain Res 96:25–40.

Isa K, Sooksawate T, Kobayashi K, Kobayashi K, Redgrave P, Isa T (2020) Dissecting the Tectal Output Channels for Orienting and Defense Responses. eneuro 7:ENEURO.0271-0220.2020.

Jo HJ, Saad ZS, Simmons WK, Milbury LA, Cox RW (2010) Mapping sources of correlation in resting state FMRI, with artifact detection and removal. Neuroimage 52:571–582.

Jun E, Na KS, Kang W, Lee J, Suk HI, Ham BJ (2020) Identifying resting-state effective connectivity abnormalities in drug-naïve major depressive disorder diagnosis via graph convolutional networks. Hum Brain Mapp 41:4997–5014.

Koller K, Rafal RD, Platt A, Mitchell ND (2019) Orienting toward threat: Contributions of a subcortical pathway transmitting retinal afferents to the amygdala via the superior colliculus and pulvinar. Neuropsychologia 128:78–86.

Kragel PA, Ceko M, Theriault J, Chen D, Satpute AB, Wald LW, Lindquist MA, Feldman Barrett L, Wager TD (2021) A human colliculus-pulvinar-amygdala pathway encodes negative emotion. Neuron 109:2404–2412 e2405.

Lin JY, Murray SO, Boynton GM (2009) Capture of attention to threatening stimuli without perceptual awareness. Curr Biol 19:1118–1122.

Mai JKM, M.; Paxinos, G. (2016) Atlas of the human brain, 4th Edition: Acadamic Press.

May PJ (2006) The mammalian superior colliculus: laminar structure and connections. Prog Brain Res 151:321–378.

Merkle EC, Rosseel Y (2018) blavaan: Bayesian Structural Equation Models via Parameter Expansion. J Stat Softw 85.

Pegna AJ, Khateb A, Lazeyras F, Seghier ML (2005) Discriminating emotional faces without primary visual cortices involves the right amygdala. Nat Neurosci 8:24–25.

Pelli DG (1997) The VideoToolbox software for visual psychophysics: transforming numbers into movies. Spat Vis 10:437–442.

Pessoa L, Adolphs R (2010) Emotion processing and the amygdala: from a ‘low road’ to ‘many roads’ of evaluating biological significance. Nat Rev Neurosci 11:773–782.

Pzxinos GH, X.; Sengul, G.; Watson, C. (2012) Organization of brainstem nuclei. In: The Human Nervous System, 3rd Edition, pp 260–327. Amsterdam: Elsevier Acaemic Press.

Rafal RD, Koller K, Bultitude JH, Mullins P, Ward R, Mitchell AS, Bell AH (2015) Connectivity between the superior colliculus and the amygdala in humans and macaque monkeys: virtual dissection with probabilistic DTI tractography. J Neurophysiol 114:1947–1962.

Rissman J, Gazzaley A, D’Esposito M (2004) Measuring functional connectivity during distinct stages of a cognitive task. Neuroimage 23:752–763.

Rosseel Y (2012) lavaan: An R Package for Structural Equation Modeling. J Stat Softw 48:1–36.

Salay LD, Ishiko N, Huberman AD (2018) A midline thalamic circuit determines reactions to visual threat. Nature 557:183–189.

Schiller PH, Stryker M, Cynader M, Berman N (1974) Response characteristics of single cells in the monkey superior colliculus following ablation or cooling of visual cortex. J Neurophysiol 37:181–194.

Shang C, Liu Z, Chen Z, Shi Y, Wang Q, Liu S, Li D, Cao P (2015) BRAIN CIRCUITS. A parvalbumin-positive excitatory visual pathway to trigger fear responses in mice. Science 348:1472–1477.

Shang C, Chen Z, Liu A, Li Y, Zhang J, Qu B, Yan F, Zhang Y, Liu W, Liu Z, Guo X, Li D, Wang Y, Cao P (2018) Divergent midbrain circuits orchestrate escape and freezing responses to looming stimuli in mice. Nat Commun 9:1232.

Sun H, Frost BJ (1998) Computation of different optical variables of looming objects in pigeon nucleus rotundus neurons. Nat Neurosci 1:296–303.

Tailby C, Cheong SK, Pietersen AN, Solomon SG, Martin PR (2012) Colour and pattern selectivity of receptive fields in superior colliculus of marmoset monkeys. J Physiol 590:4061–4077.

Tardif E, Clarke S (2002) Commissural connections of human superior colliculus. Neuroscience 111:363–372.

Thieu MK, Ayzenberg V, Lourenco SF, Kragel PA (2024) Visual looming is a primitive for human emotion. iScience 27:109886.

Tseng Y-T, Schaefke B, Wei P, Wang L (2023) Defensive responses: behaviour, the brain and the body. Nat Rev Neurosci 24:655–671.

Wall MB, Walker R, Smith AT (2009) Functional imaging of the human superior colliculus: an optimised approach. Neuroimage 47:1620–1627.

Wang Y, Frost BJ (1992) Time to collision is signalled by neurons in the nucleus rotundus of pigeons. Nature 356:236–238.

Wei P, Liu N, Zhang Z, Liu X, Tang Y, He X, Wu B, Zhou Z, Liu Y, Li J, Zhang Y, Zhou X, Xu L, Chen L, Bi G, Hu X, Xu F, Wang L (2015) Processing of visually evoked innate fear by a non-canonical thalamic pathway. Nat Commun 6:6756.

Zhang P, Zhou H, Wen W, He S (2015) Layer-specific response properties of the human lateral geniculate nucleus and superior colliculus. Neuroimage 111:159–166.

Zhang P, Wen W, Sun X, He S (2016) Selective reduction of fMRI responses to transient achromatic stimuli in the magnocellular layers of the LGN and the superficial layer of the SC of early glaucoma patients. Hum Brain Mapp 37:558–569.

Zhaoping L (2016) From the optic tectum to the primary visual cortex: migration through evolution of the saliency map for exogenous attentional guidance. Curr Opin Neurobiol 40:94–102.

Zhou Z, Liu X, Chen S, Zhang Z, Liu Y, Montardy Q, Tang Y, Wei P, Liu N, Li L, Song R, Lai J, He X, Chen C, Bi G, Feng G, Xu F, Wang L (2019) A VTA GABAergic Neural Circuit Mediates Visually Evoked Innate Defensive Responses. Neuron 103:473–488 e476.

Zhuang J, LaConte S, Peltier S, Zhang K, Hu X (2005) Connectivity exploration with structural equation modeling: an fMRI study of bimanual motor coordination. Neuroimage 25:462–470.

Zubair M, Murris SR, Isa K, Onoe H, Koshimizu Y, Kobayashi K, Vanduffel W, Isa T (2021) Divergent Whole Brain Projections from the Ventral Midbrain in Macaques. Cereb Cortex 31:2913–2931.

